# Blood-based genome-wide DNA methylation correlations across body fat and adiposity-related biochemical traits

**DOI:** 10.1101/2023.03.05.530890

**Authors:** Alesha A Hatton, Robert F Hillary, Elena Bernabeu, Daniel L McCartney, Riccardo E Marioni, Allan F McRae

## Abstract

The recent increase in obesity levels across many countries is likely to be driven by nongenetic factors. The epigenetic modification DNA methylation (DNAm) may help to explore this as it is sensitive to both genetic and environmental exposures. While the relationship between DNAm and body fat traits has been extensively studied [1–9], there is limited literature on the shared associations of DNAm variation across such traits. Akin to genetic correlation estimates, which measure the degree of common genetic control between two traits, here we introduce an approach to evaluate the similarities in DNAm associations between traits, DNAm correlations. As DNAm can be both a cause and consequence of complex traits [5, 10, 11], DNAm correlations have the potential to provide novel insights into trait relationships above that currently obtained from genetic and phenotypic correlations. Utilising 7,519 unrelated individuals from Generation Scotland (GS), we calculated DNAm correlations using the bivariate OREML framework in the OSCA software [12] to investigate the shared associations of DNAm variation between traits. For each trait we also estimated the shared contribution of DNAm between sexes. We identified strong, positive DNAm correlations between each of the body fat traits (BMI, body fat % and waist to hip ratio; ranging from 0.96 to 1.00), finding larger associations than those identified by genetic and phenotypic correlations. We identified a significant deviation from 1 in the r_DNAm_ for BMI between males and females, with sex-specific DNAm changes associated with BMI identified at eight DNAm probes. Employing genome-wide DNAm correlations to evaluate the similarities in the associations of DNAm with complex traits has provided novel insight into obesity related traits beyond that provided by genetic correlations.

## Introduction

Obesity constitutes a growing healthcare burden and is a major risk factor for several chronic diseases including cardiovascular diseases and diabetes [13, 14]. Body mass index (BMI), the most widely used measure of obesity, results from the complex interplay between genetic, environmental, and modifiable lifestyle factors. The increase in BMI levels in recent years [15] is likely to be driven by nongenetic factors. DNA methylation (DNAm) is a commonly studied epigenetic modification that is responsive to both genetics and the environment, making it an ideal target for studying the consequences of modifiable health factors, such as obesity. The relationship between DNAm and BMI, as well as other body fat and adiposity-related biochemical traits, has been extensively studied [1–9]. However, the shared associations of DNAm variation across such traits represents an important gap in our understanding of the biological processes pertaining to obesity.

Akin to genetic correlation estimates, which measure the degree of common genetic control between two traits, here, we introduce an approach to evaluate the similarities in DNAm associations between traits, DNAm correlations (r_DNAm_). In contrast to genetic variants, DNAm reflects a wide range of environmental exposures and may reflect the cumulative burden of adverse exposures throughout the life course. In addition, variation in DNAm has been implicated as arising from individual differences in traits such as BMI and smoking [5, 10], with some evidence suggesting BMI in childhood may be predictive of adolescent DNAm levels at sites throughout the genome [1]. Thus, while genetic correlations capture causal effects on the traits, DNAm correlations will capture consequence too. Ascertaining effects from both directions may result in the detection of additional biological mechanisms underlying the relationship between these traits. We also recognise that with a large portion of the DNA methylome under genetic control [11], DNAm correlations will likely capture part of the shared genetic contribution between these traits. However, recent work [4, 5, 10, 16] has demonstrated that DNAm associated with BMI trait variance is independent of genetic variation. This indicates that DNAm correlations have the potential to provide novel insights into trait relationships as well as the molecular underpinnings and subsequent consequences of these traits above that currently obtained from genetic correlations.

We estimate DNAm correlations for six body fat and adiposity-related biochemical traits for 7,519 unrelated individuals from Generation Scotland (GS). DNAm correlations are estimated by extending the OREML method in the OSCA software [12] to a bivariate model, akin to bivariate GREML as implemented in the GCTA software [17, 18]. These DNAm correlation estimates provide a measure of the shared similarity of DNAm variation between phenotypes, noting that while SNPs explain the variation in traits, DNAm only captures this variation and reflects both cause and consequence. We compared these DNAm correlations to genetic and phenotypic correlations to investigate if they provide novel insights into the molecular underpinnings of these traits.

## Results

Generation Scotland (GS) is a Scottish family-based study with over 24,000 participants recruited between 2006 and 2011 [19, 20]. We analysed data from 7,519 unrelated individuals (genetic relationship matrix (GRM) pruned at 0.05) from the larger GS dataset to avoid confounding between genetic relatedness and epigenetic similarity. Blood-based DNAm levels at 781,379 DNAm sites were quantified using the Illumina Methylation EPIC array in three sets based on time of generation of DNAm array processing. Three anthropometric measurements and three biochemical phenotypes were investigated: body mass index (BMI; kg/m^2^), body fat percentage (%), waist to hip ratio (WHR), glucose (mmol/L), high-density lipoprotein cholesterol (HDL, mmol/L) and total cholesterol (mmol/L). Demographic and summary information from Generation Scotland (GS) for the six phenotypes are presented in Table 1. We estimated the proportion of phenotypic variation captured by DNAm for each trait based on a methylation relationship matrix (MRM) using OSCA [12], the variation explained by SNPs based on a GRM, as well as that captured jointly by DNAm and SNPs. As demonstrated previously [10], we observed non-zero estimates for the proportion of variance captured by DNAm when estimated jointly with SNPs which demonstrates that some of the variation captured by DNAm is additional to that being captured by SNPs (see Supplementary Results, Supplementary Figure 1 and Supplementary Table 1). The additional variation captured by DNAm indicates that there is a potential to gain novel insights into trait relationships with DNAm correlations that are not currently captured by genetic correlations based on common SNPs.

**Table 1:**
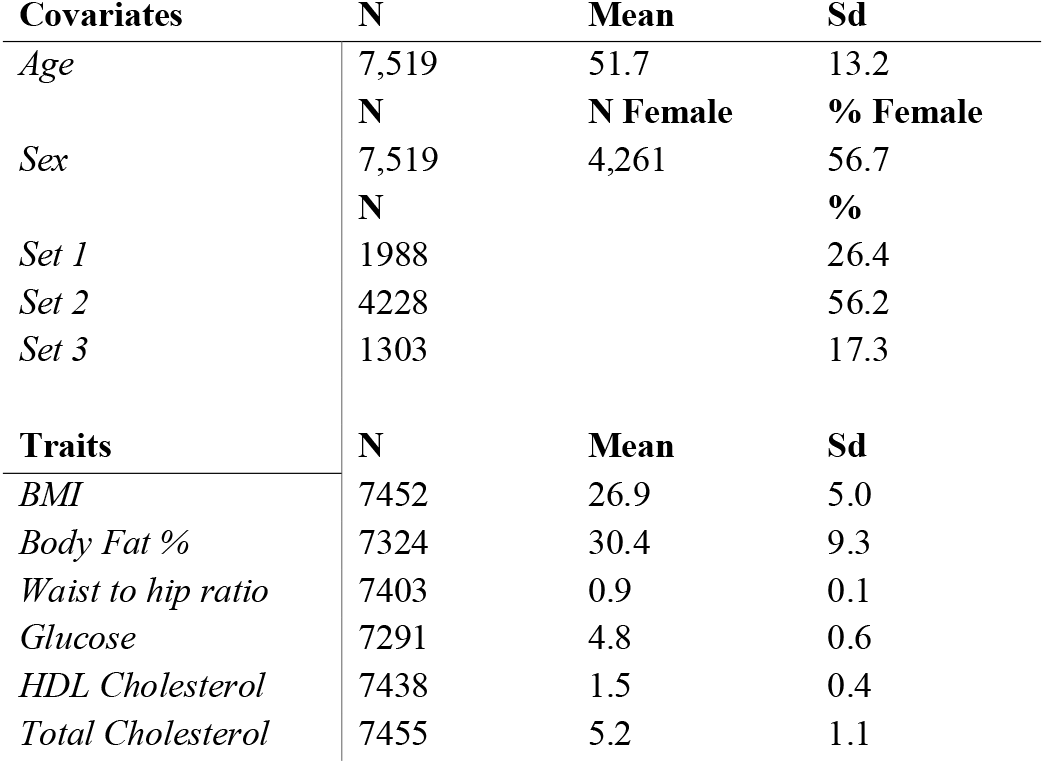
Cohort summary for Generation Scotland (GS; N=7,519)

We extended the OREML approach of the OSCA to a bivariate model that simultaneously estimates the proportion of variance in the two traits captured by DNAm as well as quantifying the shared associations between DNAm and the two traits. We term this shared association as a DNAm correlation, or r_DNAm_, reflecting the similarity of the approach to estimating genetic correlations via the GREML model.

### DNAm correlation between sets

As a proof-of-principle illustration of the application of genetic correlation methods to DNAm, and to demonstrate strong concordance between the three sample sets with GS, we estimated the DNAm correlation of the six traits across sample set. The underlying assumption is that there should be no inter-set variation in contribution of DNAm to each of the traits and thus the DNAm correlation estimates should not be different from 1. We first test this assumption by performing EWAS within each set for each trait to determine if there is concordance in probe effects using simple linear regression in the OSCA software [12]. Due to differences in sample size between the sets which impacts discovery, only those probes that were nominally significant across all sets (p<0.001) were compared. The concordance in probe association coefficients was evaluated using Pearson’s correlation and was found to be very high (0.95) between all sets (Supplementary Figure 3). This suggests that the estimated effect sizes between DNAm and each of the traits is consistent between sets. We subsequently calculated the DNAm correlation between sets for each trait. Most of the correlations were found to not significantly deviate from 1 (p<0.05) consistent with our expectation (Figure 1, Supplementary Table 4). We note that the slight deviations from 1 observed for glucose and HDL cholesterol as well as large standard errors, while not significant after adjusting for multiple testing using a Bonferroni correction, may reflect deviations from sample collection protocols and measurement errors rather than a reflection of the method.

**Figure 1:**
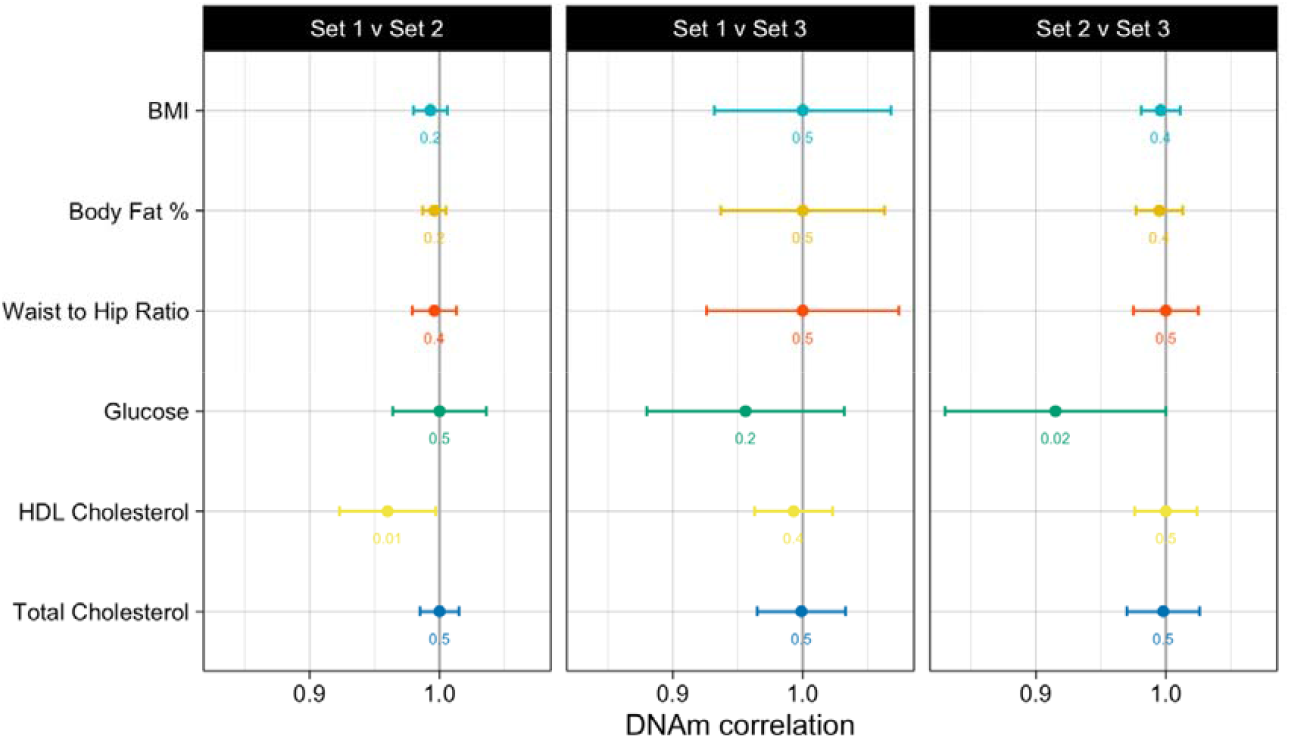
DNAm correlation between sets for each trait. The DNAm correlations between each of the set pairs is displayed on the x-axis with standard errors indicated by error bars. P-values from a log likelihood test for the hypothesis of fixing the DNAm correlation at 1 are presented in text below each estimate and in Supplementary Table 4.

### DNAm Correlation between traits

We estimated the DNAm correlation between the six body fat related phenotypes using the bivariate OREML framework that estimates the similarity of DNAm associations between traits. The DNAm correlations are presented in Figure 2 (Supplementary Table 7) alongside genetic correlations calculated using the bivariate GREML framework with a GRM implemented in the GCTA software [18] and phenotypic correlations. We identified strong, positive DNAm correlations between each of the body fat traits (BMI, body fat % and waist to hip ratio; ranging from 0.96 to 1.00), with correlation between BMI and waist to hip ratio found to be not significant different from 1 (r_DNAm_=1.00, se=0.0005). These associations were observed to be of greater magnitude than both genetic (r_G_ ranging from 0.65 to 0.86) and phenotypic correlations (r_P_ ranging from 0.51 to 0.85). The body fat traits demonstrated moderate DNAm correlations with glucose (r_DNAm_ ranging from 0.42 to 0.62), again of a greater magnitude than both genetic and phenotypic correlations. We observed negative DNAm correlations between each of the body fat traits and HDL cholesterol, with a slightly stronger correlation observed for BMI. These correlations were in the same direction as genetic correlations, with a similar magnitude while phenotypic correlations were observed to be closer to zero. DNAm correlations for each of the body fat traits with total cholesterol were found to not be significantly different from zero. This is consistent with genetic correlations between total cholesterol and both BMI and body fat %, while the genetic correlation between waist to hip ratio and total cholesterol was non zero (r_DNAm_=0.43, se=0.22, pvalue=0.02). Similarly, DNAm correlations between HDL cholesterol and glucose were observed to be similar to genetic correlations although of slightly less magnitude. We observed moderate positive DNAm correlation between total cholesterol and both glucose and HDL cholesterol while the genetic correlation was found to be not significantly different from zero between these trait pairs. Further, we demonstrate these results are independent of variance attributable to data structure, by finding practically identical estimates for DNAm correlations when adjusting for the first 20 principal components of the DNAm levels and the first 20 principal components of the genetic data (Supplementary Table 8).

**Figure 2:**
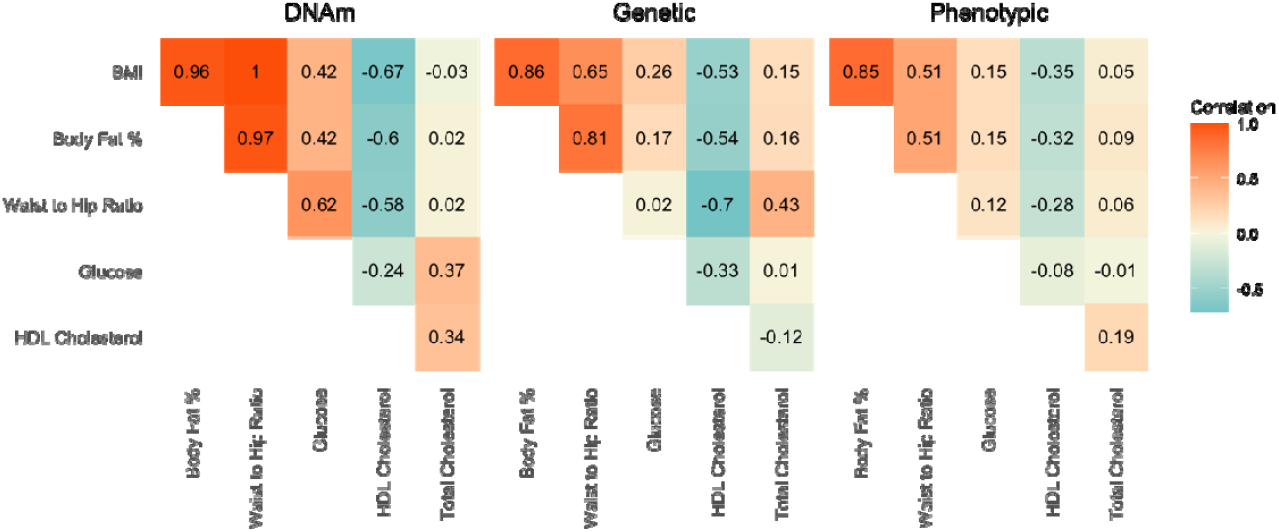
DNAm (left), genetic (middle) and phenotypic (right) correlations among six traits. Red, positive correlation; blue, negative correlation.

### DNAm correlations between sexes

Given previously reported genetic [21–23] and DNAm [4, 24] sex differences for body fat-related traits, we investigated if the contribution of DNAm for each trait was consistent across sex. First, we estimated the proportion of variance captured by DNAm in each sex. For BMI and body fat percentage the proportion of variance captured by DNAm was largely consistent across sexes while for waist to hip ratio, glucose, HDL cholesterol and total cholesterol the variance captured by DNAm in males was greater than that captured in females (Figure 3, Supplementary Table 5). Next DNAm correlations were calculated for each trait between sexes. Two traits were identified as having DNAm correlations significantly different from 1 however only BMI survived multiple testing using a Bonferroni correction (BMI r_DNAm_=0.79, se 0.07, pvalue=7.0×10^-4^; WHR r_DNAm_=0.95, se=0.04, pvalue=0.016; Figure 3 and Supplementary Table 6). Several previous studies have presented genetic correlations for BMI between the sexes that were significantly different from 1 (ranging from 0.93 to 0.96; [21–23]) however the greater deviation between the sexes captured by DNAm correlation potentially suggests the presence of novel sex differential biological consequences of BMI.

**Figure 3:**
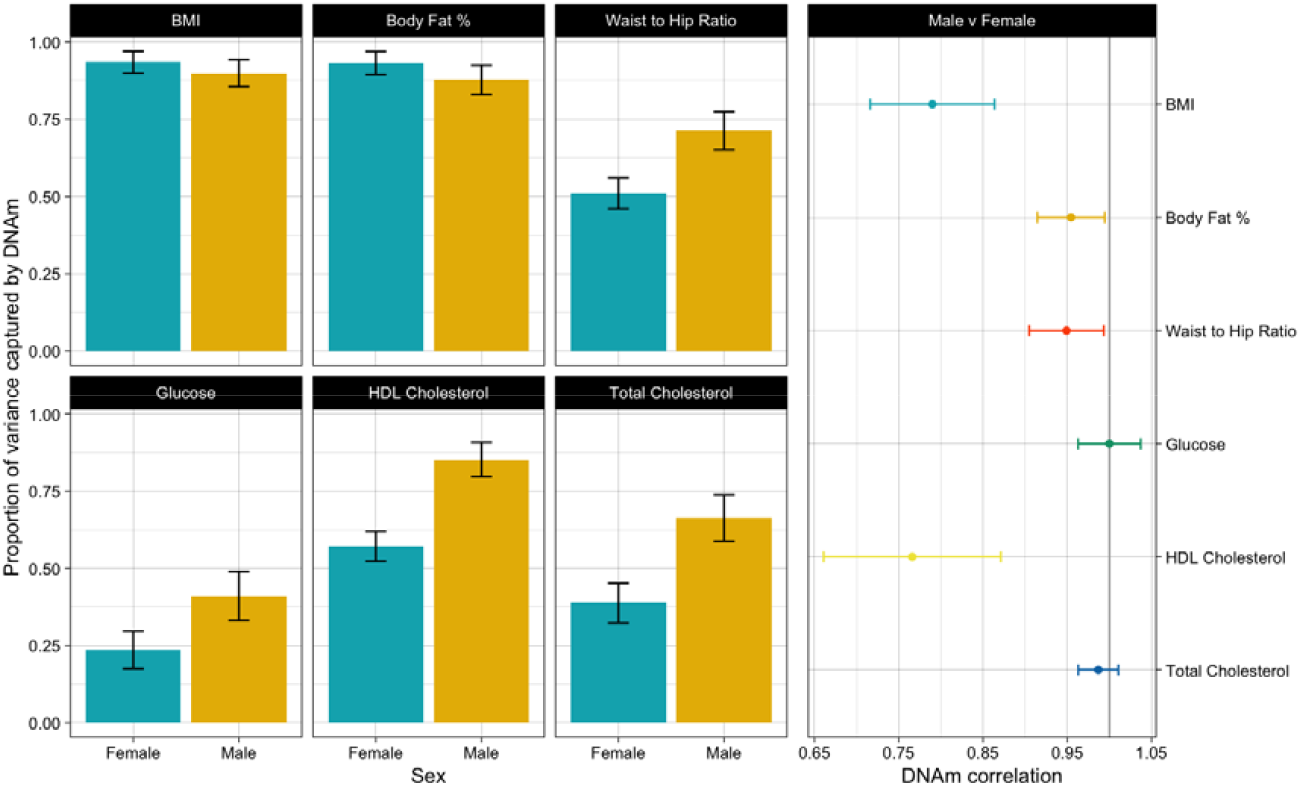
The proportion of phenotypic variance captured by DNAm by sex for each trait (left) and the DNAm correlation between sexes for each trait (right).

We further examined sex differences in the contribution of DNAm for BMI by investigating the presence of probe-by-sex interactions. We performed an EWAS for BMI including probe, sex and the interaction between probe and sex as covariates in a linear model. We identified eight probes across four chromosomes with significant probe-by-sex interactions at p<6.4×10^-8^, which is Bonferroni corrected for the number of DNAm probes analysed. We note this set of probes represented six independent probes, with two pairs of probes that were closely located together likely co-methylated (correlation between DNAm M-values>0.8 between probe pairs: cg16936953 and cg12054453, and cg18181703 and cg11047325). For all eight probes DNAm was higher for females as BMI increased, with no significant association observed in males (Table 2), with trend plots provided in Supplementary Figure 4. This may reflect a unique response in females to BMI levels.

**Table 2:** DNAm probes identified with probe by sex interactions (p<6.4×10^-8^) with BMI.

| Probe ID | CHR | Probe BP | CpG Island | Related Gene | Probe effect in Females | Probe se in Females | Probe pvalue in females | Interaction effect | Interaction se | Interaction pvalue |
| --- | --- | --- | --- | --- | --- | --- | --- | --- | --- | --- |
| cg12269535 | 6 | 43142014 | Shore | SRF | -0.66 | 0.07 | $3.6 \times 10^{-20}$ | 0.60 | 0.11 | $4.0 \times 10^{-08}$ |
| cg16936953 | 17 | 57915665 | Open sea | VMP1 | -0.37 | 0.05 | $1.3 \times 10^{-14}$ | 0.40 | 0.07 | $3.3 \times 10^{-08}$ |
| cg12054453 | 17 | 57915717 | Open sea | VMP1 | -0.27 | 0.04 | $6.4 \times 10^{-14}$ | 0.31 | 0.05 | $9.0 \times 10^{-09}$ |
| cg19748455 | 17 | 76274856 | Open sea | LOC100996291 | -0.75 | 0.06 | $1.3 \times 10^{-30}$ | 0.63 | 0.10 | $2.2 \times 10^{-10}$ |
| cg18181703 | 17 | 76354621 | Shore | SOCS3 | -0.79 | 0.07 | $6.2 \times 10^{-28}$ | 0.60 | 0.11 | $1.3 \times 10^{-08}$ |
| cg11047325 | 17 | 76354934 | Island | SOCS3 | -0.43 | 0.04 | $2.6 \times 10^{-25}$ | 0.35 | 0.06 | $1.1 \times 10^{-08}$ |
| cg00840791 | 19 | 16453259 | Intergenic | | -0.45 | 0.03 | $8.6 \times 10^{-47}$ | 0.29 | 0.05 | $7.4 \times 10^{-10}$ |
| cg09349128 | 22 | 50327986 | Shore | CRELD2 | -1.05 | 0.08 | $2.5 \times 10^{-39}$ | 0.79 | 0.12 | $1.3 \times 10^{-10}$ |

## Discussion

We investigated the shared associations of DNAm variation between body fat and adiposity-related biochemical traits by extending the OREML framework to a bivariate model, similar to the estimation of genetic correlations through GREML. For the majority of trait pairs the DNAm correlations, whilst strongly concordant in direction, were observed to be greater in magnitude compared to both genetic and phenotypic correlations, particularly between body fat traits. There are several potential explanations for this. DNAm is known to capture risk factors beyond genetics, suggesting DNAm correlations are likely capturing common environmental or lifestyle factors between traits such as dietary factors. DNAm correlation may also be capturing common consequence of these traits, that is the consequence of both traits affecting downstream pathways e.g. inflammation. This hypothesis is supported by previous studies which have demonstrated that while large amounts of the phenotypic variance can be captured by DNAm for some traits (e.g. BMI and smoking), for the most part these have been implicated as arising from trait consequence [5, 10]. In particular, Wahl et al. suggested that changes in DNAm (measured in blood and adiposity tissue) associated with BMI may be the consequence of changes in lipid and glucose metabolism associated with BMI [5]. This ascertainment of both causal and consequential effects may explain why DNAm correlations were observed to be of greater magnitude than their genetic counterparts. The strong positive DNAm correlations between each of the body fat traits is consistent with DNAm derived from whole blood reflecting a general response to adiposity, while genetic correlations are capturing differences in the genetic control of specific fat distribution. Support for such a conclusion in literature is conflicting. A recent study of DNAm in adipose tissue in women identified associations with body fat distribution, of which 50% of sites replicated whole blood derived DNAm [25]. Several other studies have demonstrated strong overlap between CpG sites associated with BMI, waist circumference and body fat % indicating common methylation sites are similarly influenced by both general and abdominal obesity [26–28]. However, Crocker *et al*. [28] found a low degree of overlap between waist circumference and body fat percentage from subsequent gene ontology enrichment and differentially methylated region analyses, suggesting these measurements represent biologically distinct concepts. We note the inconsistency in conclusions from Crocker *et al*. may have been impacted by the investigation of overlap in significant results rather than formally testing for differences and additionally limited by sample size (N=2,325).

We also recognise that, given a large portion of the DNA methylome is under genetic control [11], DNAm correlations are likely capturing part of the shared genetic contribution between these traits. We demonstrated that a large portion of the phenotypic variation captured by DNAm is separate from that being explained by SNPs, a conclusion which is supported in the literature [4, 5, 10, 16]. Further, we identified independence between the MRM and GRM when fit as random effects in the univariate GREML framework using the CORE-GREML approach. Despite this, we were unable to formally determine if the contribution of DNAm which was shared between traits was similarly separate of the shared genetic influence. This limitation in our study was likely due to sample size, with joint MRM and GRM bivariate REML models unable to converge and therefore unable to estimate DNAm correlations conditional on SNPs.

Given previously reported genetic [21–23] and DNAm [4, 24] sex differences for body fat related traits, we investigated whether these are also captured by DNAm correlations. We identified a significant deviation from 1 in the DNAm correlation for BMI between males and females. Several previous studies have presented genetic correlations for BMI between the sexes that were significantly different from 1 (ranging from 0.93 to 0.96; [21–23]). The greater deviation captured by the DNAm correlation however potentially suggests the presence of sex differential novel biological consequences of BMI. We further identified eight DNAm probe-by-sex interactions for BMI (which represent six independent DNAm sites), observing hypermethylation in females as BMI increased, with no associated observed in males. Of note, all but one of these probes having been previously shown to be associated with BMI [4, 5, 27, 29, 30]. In particular, probe cg18181703 is located the *SOCS3* gene, a suppressor of the cytokine signalling pathway, and has been found to be inversely associated with BMI, waist to hip ratio, triglycerides and metabolic syndrome, and positively associated with HDL [4]. It has also been shown to moderate the effect of cumulative stress on obesity [31]. DNAm of cg09349128 located in *PIM3*, a gene involved in energy metabolism, has been found to mediate the association between famine exposure and BMI. Additionally, probes cg16936953 and cg12054453 are located in the *VMP1* gene, which has been implicated broadly in lipid homeostasis and regulation in the formation of lipid droplets and lipoproteins, for which dysregulation is involved in a variety of diseases including obesity, fatty liver disease and cholesterol ester storage [32, 33]. We also found evidence for a probe-by-sex interaction with DNAm at probe cg12269535 located in the *SRF* gene, which is associated with insulin resistance and may contribute to the pathogenesis of Type 2 Diabetes [34]. We note that probe-by-sex interactions have been previously investigated in the context of BMI [4, 35], with each study identifying only a single CpG, however we were unable to replicate any previous findings (Supplementary Figure 5).

We recognise there are some caveats and further considerations for this work. The EPIC array captures only a small proportion of the methylome, with Hillary et al. previously demonstrating that decreasing the number of methylation sites reduces estimates of variance captured by DNAm and prediction metrics [36]. This impacts the interpretability our analyses as a low variance captured by DNAm doesn’t necessarily indicate a lack of correlation between DNAm and traits as DNAm sites which are unmeasured may contribute to the association. As such, greater coverage may resultingly influence DNAm correlation estimates. Similarly, while variance component estimation based on DNAm requires smaller samples sizes than needed for accurate estimation of genetic correlation due to the MRM capturing more variance, there is value in increasing sample sizes as well. In these analyses we were unable to report on DNAm correlations conditional on SNPs as joint MRM and GRM bivariate REML models were unable to convergence. We attempt to address this by adjusting univariate and bivariate OREML models based on DNAm with covariate adjustment for first 20 principal components of the genetic data. We find models with and without these adjustments yield practically identical estimates for both proportion of variance captured and DNAm correlations. While it has been previously shown that much of the genetic control of DNAm is shared across populations [37–40], as DNAm is also responsive to the environment, it would not be unexpected for such estimates to vary by ancestry, or geography. While we suspect our results will be generalisable across comparable samples, replication in similar populations as well as populations of different ancestry, ethnicity or geography would provide greater insight into these results.

Overall, we present an approach to investigating shared biology across traits using DNAm correlations. This has provided novel insight into obesity related traits, showing the shared associations of DNAm between BMI, waist to hip ratio and body fat %, beyond that recognised through genetic correlation analysis and has identified sex specific DNAm changes associated with BMI.

## Acknowledgements

All components of GS received ethical approval from the NHS Tayside Committee on Medical Research Ethics (REC Reference Number: 05/S1401/89). GS has also been granted Research Tissue Bank status by the East of Scotland Research Ethics Service (REC Reference Number: 20-ES-0021), providing generic ethical approval for a wide range of uses within medical research. GS received core support from the Chief Scientist Office of the Scottish Government Health Directorates (CZD/16/6) and the Scottish Funding Council (HR03006). Genotyping and DNA methylation profiling of the GS samples was carried out by the Genetics Core Laboratory at the Edinburgh Clinical Research Facility, Edinburgh, Scotland, and was funded by the Medical Research Council UK and the Wellcome Trust (Wellcome Trust Strategic Award Stratifying Resilience and Depression Longitudinally (STRADL; Reference 104036/Z/14/Z). The DNA methylation data assayed for Generation Scotland was partially funded by a 2018 NARSAD Young Investigator Grant from the Brain & Behavior Research Foundation (Ref: 27404; awardee: Dr David M Howard) and by a JMAS SIM fellowship from the Royal College of Physicians of Edinburgh (Awardee: Dr Heather C Whalley). AAH is supported by an Australian Government Research Training Program (RTP) Scholarship. AFM is supported by an Australian Research Council Future Fellowship (FT200100837).

## Methods

### Study cohort

All data for the study came from Generation Scotland: Scottish Family Health Study (GS). The family-based genetic epidemiological cohort consists of over 24,000 volunteers which has been described previously [19, 20]. Recruitment took place between 2006 and 2011, when individuals their family members aged 18+ years were invited to a baseline clinic visit that included health questionnaires and sample donation for genomic analyses. This study uses phenotypic, DNAm and genetic data from unrelated samples (N = 7,519, GRM<0.05), with DNAm levels quantified in three sets based on time of DNAm array processing.

### Phenotypic data

Three anthropometric measurements and three biochemical phenotypes were investigated: body mass index (BMI; kg/m^2^), body fat percentage (%), waist to hip ratio, glucose (mmol/L), high-density lipoprotein (HDL) cholesterol (mmol/L) and total cholesterol (mmol/L). All phenotypes were trimmed for outliers (values that were ± 4 SDs from the mean). In addition, BMI was trimmed for extreme values at <17 and >50 kg/m2. For each trait we stratified the samples by sex then adjusted the phenotype for age and standardized the residuals by rank based inverse normal transformation before recombining the data. There was no adjustment for set as there were minimal differences in the mean across sets. Residualised phenotypes were entered as dependent variables in the subsequent analysis. Smoking pack years were calculated by multiplying the number of packs of cigarettes smoked per day by the number of years the individual has smoked and used in the adjustment of DNAm data.

### Genetic data

Genome wide genotypic details have been described previously [41]. Briefly, GS participants were genotyped with either Illumina HumanOmniExpressExome8v1-2_A or HumanOmniExpressExome-8v1_A arrays. SNPs were excluded for missing genotype call rate (>2%), and marked departure from Hardy–Weinberg equilibrium (HWE; p□<□1□×□10^-6^), low MAF (<1%). Duplicate samples were removed alongside individuals with gender mismatch and missing genotype call rate (>2%). Principal components were calculated in the GCTA software [18] using the 1,092 individuals of the 1000 Genomes population [42]. Outliers were defined as observations more than six standard deviations away from the mean of the GS individuals for the first two principal components and were subsequently removed [43]. Genotype data was imputed against HRC panel v1.1 [44]. Unrelated individuals were retained (GRM<□0.05) using the GCTA software [18]. All subsequent analyses were conducted on the unrelated individuals using HapMap3 SNPs only.

### DNA methylation data

Genome-wide blood-based DNA methylation profiled using the Illumina Methylation EPIC array and was processed in three separate sets. DNAm quality control was performed as reported previously [45]. Briefly, outliers were excluded based on the visual inspection of methylated to unmethylated log intensities, in addition to poorly performing probes and samples, and sex mismatches. Further filtering was performed to exclude non-autosomal CpG sites, CpGs that were predicted to cross-hybridise and those with polymorphisms at the target site which can alter probe binding [46, 47]. Poor performing probes, X/Y chromosome probes and participants with unreliable self-report data, saliva samples and potential XXY genotype were excluded along with probes with almost invariable beta values across individuals (standard deviation <□0.02). All 3 sets were normalised together with the final discovery dataset comprised M values at 781,379 loci for 7,519 participants. Before analysis, DNAm was adjusted in the OSCA software for age, sex, batch, slide, cell type proportions (estimated using the algorithm proposed by Houseman et al. [48]), smoking status and pack years.

### Variance component analyses

Utilising 7,519 unrelated individuals from Generation Scotland (GS), we estimate the proportion of phenotypic variance captured by genome-wide DNAm across six body fat and adiposity-related biochemical traits using omics-restricted maximum likelihood (OREML) framework in the OSCA software. This method estimates the variance captured by DNAm by construction of a DNAm relationship matrix (MRM) based on all DNAm probes and which is used to model the covariance between individuals in a univariate linear mixed model via restricted maximum likelihood (REML). This allows us to obtain the proportion of variation for each trait captured by all probes which is analogous to that of estimating SNP-based heritability based on genetic data [18, 49]. Unlike SNP-based heritability, we note that the proportion of captured by all probes may be capturing both cause and consequence of the phenotype. The GCTA software was used to calculate the GRM and similarly implemented in the OSCA software to estimate SNP-based heritability, referred to here as the proportion of phenotypic variance explained by all SNPs. We also estimated these quantities jointly in the OSCA software using --multi-orm which allows for multiple random effects.

### DNAm correlations

We estimate the DNAm correlation between phenotypes implemented using the bivariate OREML framework in OSCA utilising DNAm relationship matrices rather than the standard GRM, where the DNAm correlation is estimated from the one of the covariance components. Here the phenotypic and DNAm information came from the same unrelated individuals. This approach estimates the shared contribution of DNAm based on the MRM between phenotypes. Likelihood ratio tests were performed to test the hypotheses of fixing the correlations at both zero and one. We additionally estimated genetic correlations using GCTA and phenotypic correlations using Pearson’s correlation and compared these with r_DNAm_ to investigate if this metric provides novel insights into the molecular underpinnings of these traits. Joint estimation of rG and r_DNAm_ was not reported due as the models were unable to converge.

