## Supplementary Material for "Blood-based genome-wide DNA methylation correlations across body fat and adiposity-related biochemical traits"

**Supplementary Results**

### Estimating the proportion of variance captured by DNAm

We estimated the proportion of phenotypic variation captured by DNAm for each trait based on a methylation relationship matrix (MRM) using the OSCA software [12] as well as the variation explained by SNPs based on a GRM. We contrast these marginal estimates with those jointly calculated from both the MRM and GRM (DNAm and SNPs). These estimates allow us to determine the amount of trait variation jointly being captured by DNAm and SNPs as well as that captured by DNAm which is not explained by SNPs. The proportion of variance captured marginally by DNAm ranged from 0.29 (se 0.04) for glucose to 0.61 (0.02) for body fat percentage (Figure 1, Supplementary Table 1), with an average of 0.45 across the six traits. These estimates were consistently larger than that explained by SNPs (0.12 – 0.34) with the caveat that SNPs reflect causality while trait variation captured by DNAm may also be the result of trait consequence. We observed a slight decrease in the variance estimates for SNPs when calculated jointly with DNAm, suggesting a small overlap in the proportion of phenotypic variance being captured by the MRM and GRM (mean overlap = 0.07, range 0.02-0.1). As a proportion of the methylome is under genetic control [11] such a result is expected. We note that the decrease in SNP variance, as opposed to DNAm variance, when modelled jointly is likely a consequence of the variance estimation procedure and does not denote causality. The presence of non-zero estimates for the proportion of variance captured by DNAm when estimated jointly with SNPs demonstrates that some of the variation captured by DNAm is additional to that being captured by SNPs. This observation does not necessarily indicate the variation captured by DNAm is fully independent of that explained by genetics as not all genetic influence is captured with common SNPs their additive effects. However, the additional information captured by DNAm indicates that there is a potential to gain novel insights into trait relationships with DNAm correlations that are not currently captured by genetic correlations based on common SNPs.

When estimating variance components, GREML assumes independence between random effects. As much of the DNA methylome is under genetic control [11] and GWAS loci have been found to overlap with mQTLs for complex traits [21-23], we sought to verify if this assumption was valid in the context of phenotypic variance partitioning between MRM and GRM. A non-zero covariance between such terms can result in biased partitioning of phenotypic variance. We first investigated the relationship between the MRM and GRM by calculating the Pearson’s correlation between the off-diagonal elements of each matrix, excluding the diagonal elements as these are 1 in both matrices. We observe no correlation ($\rho$=0.004 (se 0.0002); Supplementary Figure 2). Further, we computed the Cholesky decomposition of MRM and GRM matrices to estimate the covariance between these terms using CORE GREML [24], a method derived to explicitly estimate the covariance between random effects in the GREML framework. To determine if the covariance between the MRM and GRM was non-zero, we compared CORE GREML (with covariance term) and GREML (without covariance term) using the likelihood ratio test with one degree of freedom. This was performed for each trait in the univariate GREML framework. For all traits the covariance term was non-significant (BMI p_LRT_ > 0.05, Supplementary Table 2). This suggests the assumption of independence between random effects is valid. In addition, we demonstrate the estimated the proportion of phenotypic variation captured by DNAm for each trait is independent of variance attributable to data structure by performing sensitivity analysis with covariate adjustment for shared family effects, the first 20 principal components of the DNAm levels and the first 20 principal components of the genetic data. We find that models with and without these adjustments yield practically identical estimates for the proportion of variance captured by DNAm for each trait (Supplementary Table 3).

**Supplementary Tables**

Supplementary Table 1: The proportion of variance captured for each trait. Variance components for DNAm (MRM) and SNPs (GRM) were estimated both marginally and jointly. The proportion of phenotypic variance captured for each trait (Variance) for each model is provided alongside the associated standard error (SE), pvalue (Pval) and sample size (N).

Supplementary Table 2: Likelihood ratio test comparing CORE GREML and GREML to estimate the covariance between MRM and GRM. This was performed for each trait in the univariate GREML framework.

Supplementary Table 3: Sensitivity analyses for the univariate GREML model for each trait. Sensitivity analyses were performed with covariate adjustment for the first 20 principal components of the DNAm levels (DNAm PC adjustment) and the first 20 principal components of the genetic data (SNP PC adjustment). For each model the proportion of variance captured by DNAm for each trait is presented (Variance), alongside the associated standard error (SE) and pvalue (Pval).

Supplementary Table 4: DNAm Correlation estimates by set for each trait. DNAm correlations (Correlation) are presented, alongside the associated standard error (SE) and sample size (N). Pvalues from the LRT are presented for the hypothesis of fixing the DNAm correlation at both 1 (Pval_1) and 0 (Pval_0). The proportion of variance in each trait for each set captured by DNAm as determined from the bivariate GREML model is also presented (V(O)/Vp_tr1, V(O)/Vp_tr2) as well as associated standard errors (SE).

Supplementary Table 5: The proportion of variance captured by DNAm for each trait by sex (Female and Male). The proportion of phenotypic variance captured for each trait (Variance) for each sex is provided alongside the associated standard error (SE), pvalue (Pval) and sample size (N).

Supplementary Table 6: DNAm Correlation estimates by sex for each trait. DNAm correlations (Correlation) are presented, alongside the associated standard error (SE) and sample size (N). Pvalues from the LRT are presented for the hypothesis of fixing the DNAm correlation at both 1 (Pval_1) and 0 (Pval_0). The proportion of variance in each trait for each set captured by DNAm as determined from the bivariate GREML model is also presented (V(O)/Vp_tr1, V(O)/Vp_tr2) as well as associated standard errors (SE).

Supplementary Table 7: Correlation estimates between traits calculated using each, DNAm, genetics and phenotypes. For each of the correlation measures, correlations (Correlation) are presented, alongside the associated standard error (SE) and sample size (N). For DNAm and genetic correlations, Pvalues from the LRT are presented for the hypothesis of fixing the DNAm correlation at both 1 (Pval_1) and 0 (Pval_0). The proportion of variance in each trait for each set captured by DNAm as determined from the bivariate GREML model is also presented (V(O)/Vp_tr1, V(O)/Vp_tr2) as well as associated standard errors (SE). For phenotypic correlations, the pvalue form Pearson’s correlation is presented (Pval).

Supplementary Table 8: Sensitivity analyses for the bivariate GREML model for each trait. Sensitivity analyses were performed with covariate adjustment for the first 20 principal components of the DNAm levels (DNAm PC adjustment) and the first 20 principal components of the genetic data (SNP PC adjustment). For each model, DNAm correlations (Correlation) are presented, alongside the associated standard error (SE) and sample size (N). Pvalues from the LRT are presented for the hypothesis of fixing the DNAm correlation at both 1 (Pval_1) and 0 (Pval_0). The proportion of variance in each trait for each set captured by DNAm as determined from the bivariate GREML model is also presented (V(O)/Vp_tr1, V(O)/Vp_tr2) as well as associated standard errors (SE).

**Supplementary Figures**

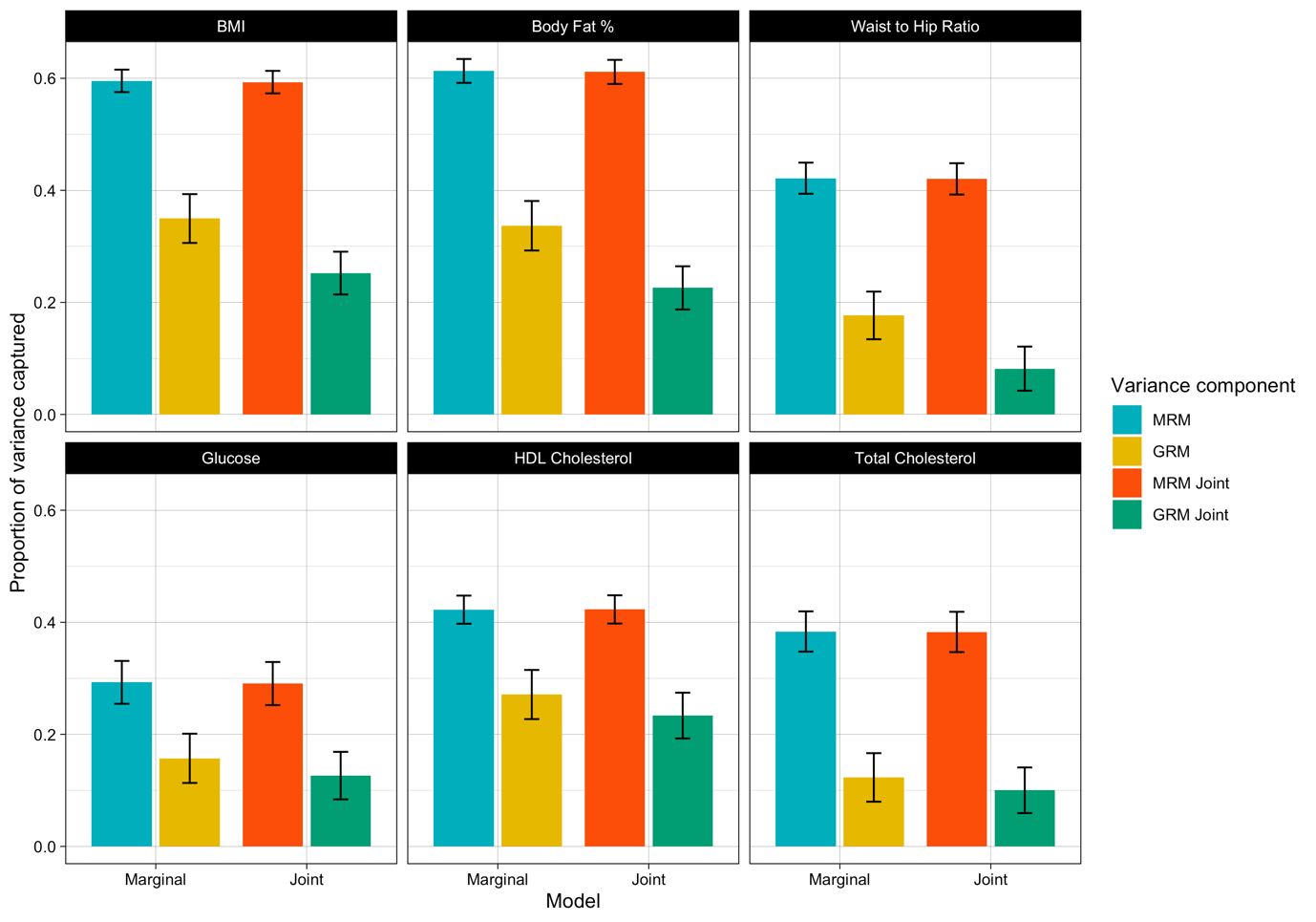

Supplementary figure 1: The proportion of variance captured for each trait. Variance components for DNAm (based on MRM) and SNPs (based on GRM) were estimated both marginally (DNAm blue, SNPs yellow) and jointly (DNAm red, SNPs green). The proportion of phenotypic variance captured for each trait for each model is displayed on the x-axis with standard errors indicated by error bars.

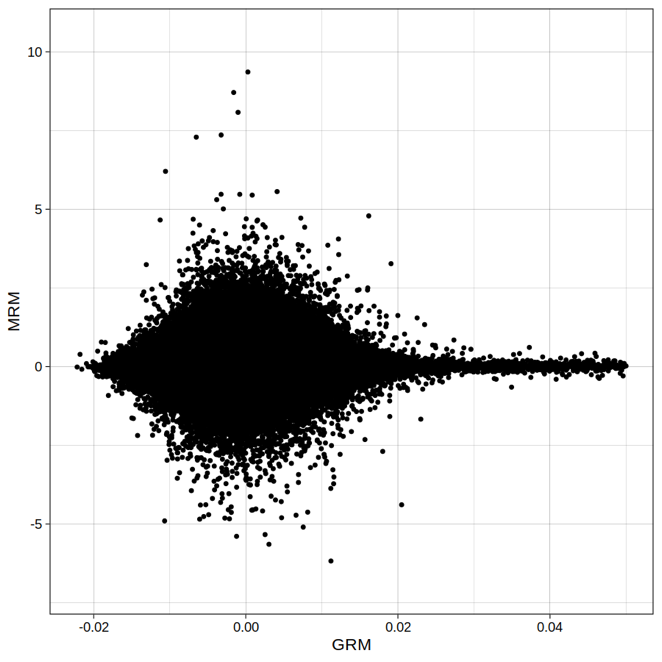

Supplementary Figure 2: Scatter plot of the off-diagonal elements of the GRM on the x-axis and MRM on the y-axis. The correlation between off-diagonal elements of GRM and MRM was 0.004 (se 0.0002).

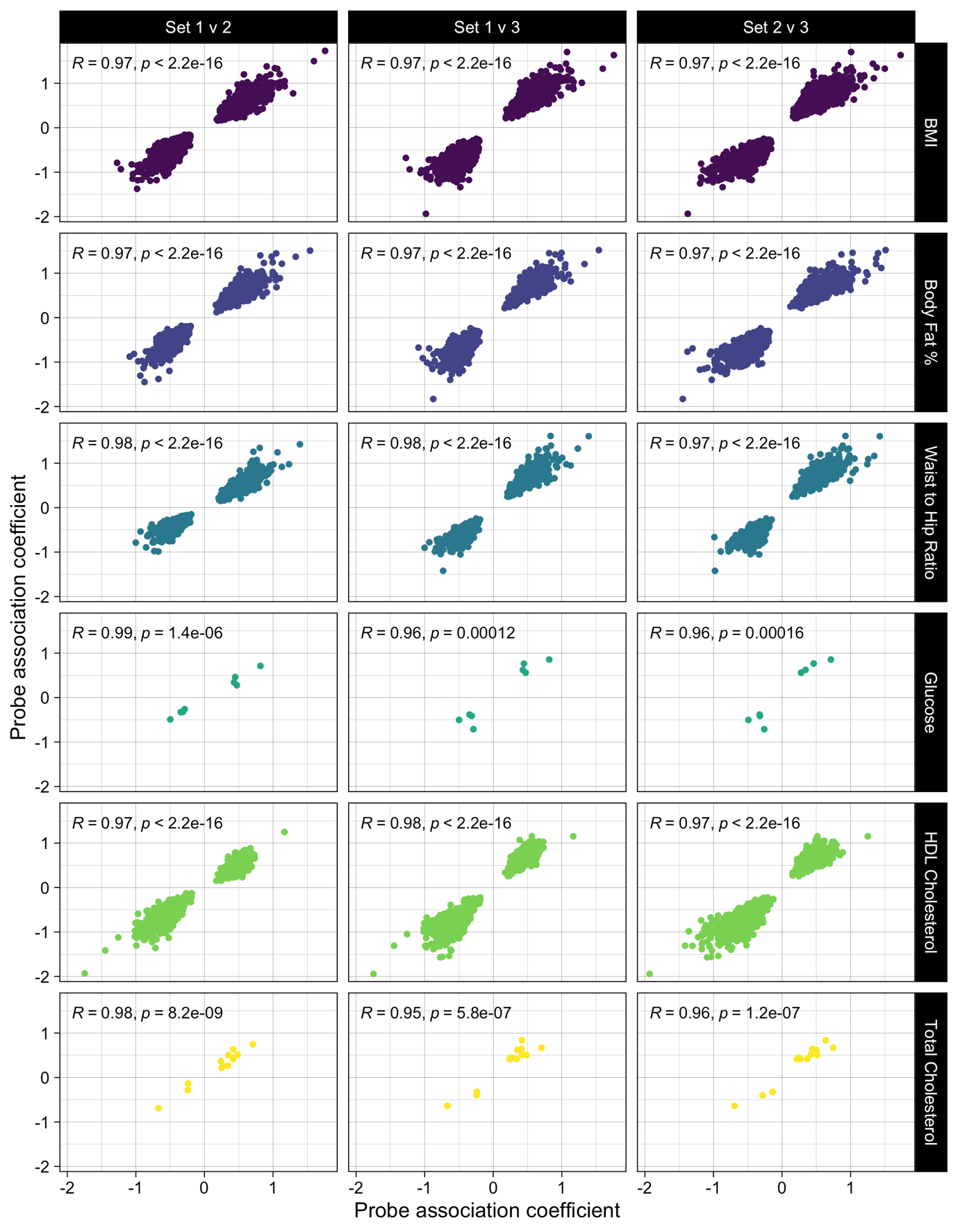

Supplementary Figure 3: Concordance of probe association coefficients from EWAS between sets for each trait. Scatter plots include probe effects for those probes that were nominally significant across all sets (p < 0.001). The concordance in probe effects was evaluated using Pearson’s correlation and was found to be high ($\rho\geq$0.95) between all sets.

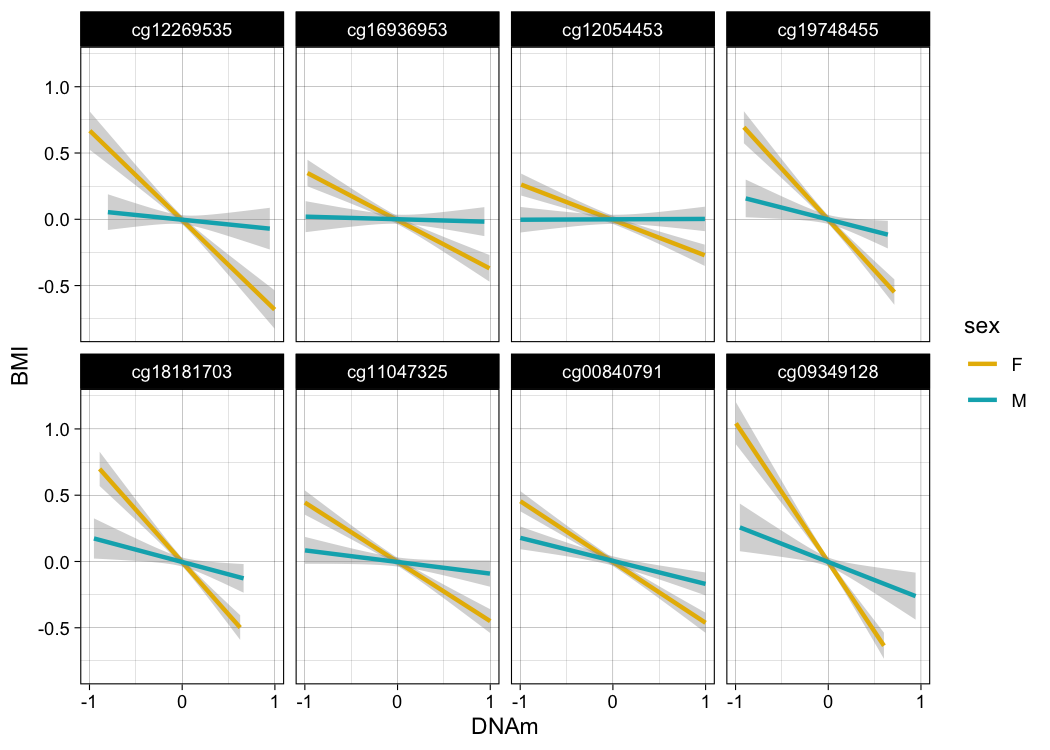

Supplementary Figure 4: Probe by sex interactions observed for BMI. The linear regression slopes for males (blue) and females (yellow) are presented for each of the eight probes with significant probe by sex interactions with BMI.

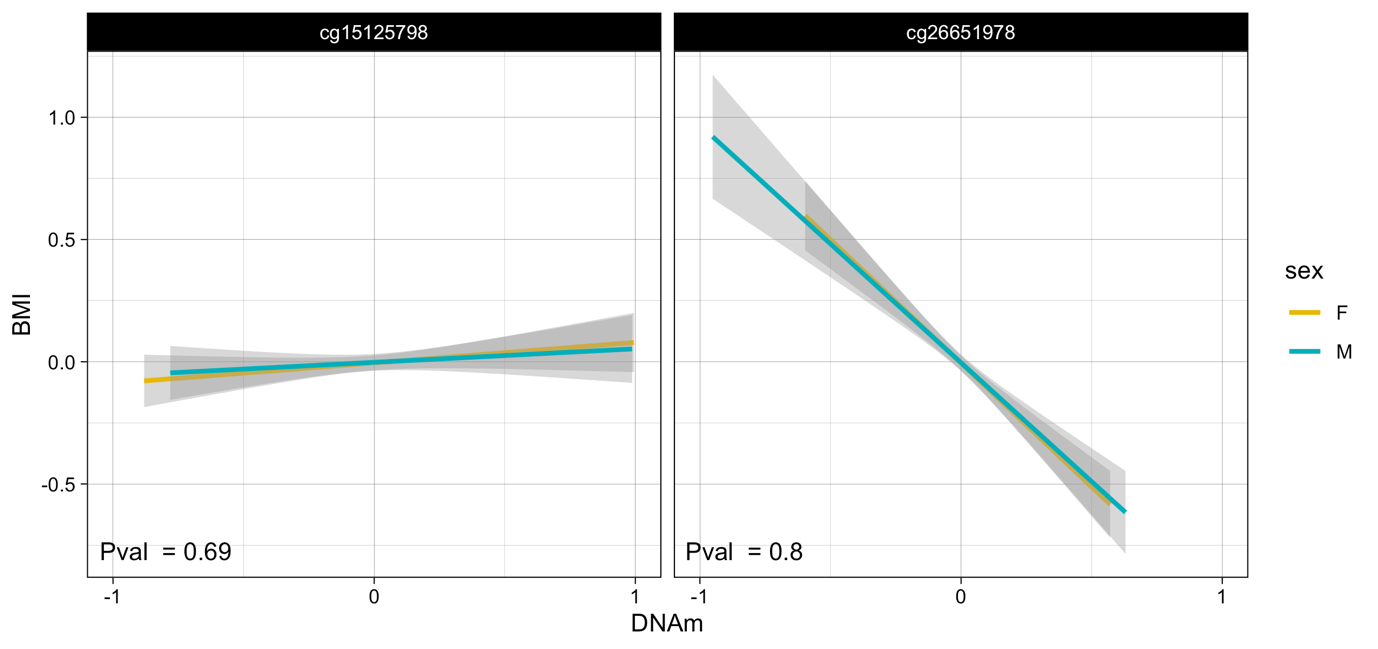

Supplementary Figure 5: Probe by sex interactions for BMI at probes cg15125798 and cg26651978. The linear regression slopes for males (blue) and females (yellow) in GS are presented for two probes previously reported in literature along each with the probe by sex interaction pvalue.
